# Battery-powered Wearable Utilizing Flexible Printed Circuit-based Organic Electrochemical Transistor Embedded with Simple Circuits of Voltage Divider and Regulator for Biosignal Measurement

**DOI:** 10.1101/2025.01.26.630991

**Authors:** Sin Yu Yeung, Haosi Lin, Yue Li, Cheuk Wang Fung, Hnin Yin Yin Nyein, I-Ming Hsing

## Abstract

With the distinctive advantages of high transconductance, low operating voltage, mixed ionic-electronic conductivities, and dynamic versatility, organic electrochemical transistor (OECT) has emerged as a promising wearable technology capable of measuring various biophysical and biochemical signals. Despite the intensive research efforts towards enhancing its wearability, challenges related to signal conversion, voltage sourcing, and manufacturing scalability are seldom addressed. Herein, we report a compact and easy-to-build integrated module that provides stable biasing from batteries while enabling current-to-voltage conversion and additional amplification of OECT’s responses. Given the known amplitude of target signals, transistor bias and amplification gain can be adjusted easily on site by tuning two key resistance values and ensuring sufficient battery voltage. Furthermore, the flexible OECTs in this work were fabricated through an industrial manufacturing process for flexible printed circuits (FPC), in which the polymeric channel material and device architecture were both customized to accommodate the fabrication constraints. Notably, preliminary measurements based on the battery-powered unit comprising our OECT and module demonstrate significantly amplified bio-signals compared to electrodes. The successful acquisition of on-body electrocardiogram voltages further underscores the potential of this platform to support current and future OECT interfaces.

## 1. Introduction

The ongoing evolution of wearable technologies is shifting their purpose from daily fitness tracking to health monitoring. Beyond measuring general physiological indicators, these technologies have been exploited for continuous monitoring of broader health metrics with improved sensitivity.^[1,2]^ Recent breakthroughs are not limited to skin-compatible integrated patch for neonatal monitoring, multimodal sweat sensors for athletic or nutritional uses, highly stretchable and adhesive 12-lead ECG Holter for detection of silent heart diseases, and high-performance pressure sensors based on eco-friendly piezoelectric materials.^[3–8]^ Together, these innovations showcase the immense, transformative potential of wearable electronics in revolutionizing contemporary healthcare and medicine.

Organic electrochemical transistor (OECT) emerges as an alternative modality that adds remarkable attributes to wearables. OECT is a three-terminal device with its active channel typically composed of poly(3,4-ethylenedioxythiophene) doped with polystyrene sulfonate (PEDOT:PSS).^[9]^ The biocompatible PEDOT:PSS-based OECT channel can directly serve as an active sensing layer, enabling simultaneous signal transduction and amplification at the sensing point.^[10–12]^ Such local amplification is favorable as it minimizes noise interference from downstream circuits.^[13]^ Additionally, PEDOT:PSS, capable of ion uptake, can achieve volumetric capacitance substantially higher than the double-layer capacitance of organic field-effect transistors (i.e. 500 versus 10 µF).^[14,15]^ This results in the highest transconductance exhibited by PEDOT:PSS-based OECT among electrolyte-gated transistors, making it exceptionally efficient in amplifying small bio-signals at low frequencies (e.g. electrocardiogram, ECG) ^[16,17]^. This property allows for an additional advantage of low operating voltage, thus reducing power consumption for wearable settings.^[18,19]^ The solution processability of PEDOT:PSS further introduces a degree of freedom in optimizing its electrical and material properties as well as transistor architecture and fabrication methods to fulfill target applications.^[20–22]^ In addition to bioelectrical measurements, OECT can effectively capture detection events of biomolecules based on its mixed ionic-to-electronic conduction, promising the integration of biochemical sensing into wearable devices.^[23,24]^

The merits of OECT specific to wearable applications have already been extensively demonstrated. Application-wise, OECT showed excellent sensitivity over electrodes in measuring the concentration of hydrogen peroxide (H_2_O_2_), an intermediate product in glucose sensing.^[25]^ When integrated into rationally designed circuits, the sensitivity and specificity of OECT in epileptic detection even surpass conventional approaches.^[26]^ It has also been reported to amplify on-body electrophysiological signals by folds of amplitude or be employed in a format of multi-channel array for human heart potential mapping.^[27,28]^ On the other hand, efforts have been invested in addressing material concerns on OECT’s wearability.^[29]^ Wearable OECTs with softness, adhesiveness, flexibility or even stretchability were developed to enhance skin compliancy and signal stability.^[30–39]^ Device longevity for long-term measurements was further accomplished by combining flexibility with anti-fouling function.^[40]^

While enhancing the properties of wearable OECT interfaces can improve measurement performance, other aspects including signal acquisition, power supply, and fabrication scalability should also be considered when developing OECT for truly wearable scenarios. For instance, the amplified output signals from OECT are in the form of current and need to be efficiently converted into voltage, the common input form for signal processing and transmission. Although current signals can be easily acquired and directly converted into digital data using a benchtop multimeter or sourcemeter, conversion methods in wireless environment require design considerations and are seldom explored.^[41–43]^ Transimpedance amplifier is one option reported to convert and actively amplify the drain-source current (*I_DS_*) of OECT without fluctuating its drain-source bias (*V_DS_*).^[25,44]^ Nonetheless, this method draws extra power due to active amplification and necessitates sophisticated circuit design. A more straightforward way is to capture output signals by floating *V_DS_* with a resistor in series, where feasibility of conversion and pre-amplification has been demonstrated given proper circuit characterizations.^[13,45]^ Regarding voltage supplies, they should be constant and adjustable to a certain extent for the OECT to maintain stability under its optimal working regime. In this case, the use of portable power sources like batteries or organic solar cells for direct OECT biasing is not recommended as this may cause fluctuations or even depreciation in its measurement performance.^[13,46–48]^ Last but not the least, the commercialization potential of wearable OECT is still determined by whether the developed protocols can be standardized into common industrial processes. Optimizing OECT fabrication from the widely adopted manufacturing processes can therefore be a good approach to achieve the captioned.

Here, we introduce a simple and easy-to-build integrated module designed for wearable OECT sensing and demonstrate its successful implementation in battery-powered wearable ECG measurements using our flexible OECTs fabricated through an industrializable process (**Figure 1**). Our developed unit is a strategic combination of voltage-divider and voltage-regulating circuits that can control signal conversion and amplification while providing stable and tunable voltages to both OECT and the circuit from batteries. The flexible devices were fabricated using the manufacturing process established for flexible printed circuits (FPC), whose compatibility is achieved via tailoring the PEDOT:PSS formulation and device architecture. Finally, our integrated system can be regulated to amplify electrophysiological signals to a higher magnitude than electrodes and is estimated to be able to operate for days before battery recharging. Overall, this work reinforces the significance of OECT in advancing wearable electronics and facilitates the realization of OECT-based wearables through the comprehensive explication of our electronic platform capable of supporting present and future OECT interfaces.

**Figure 1.**
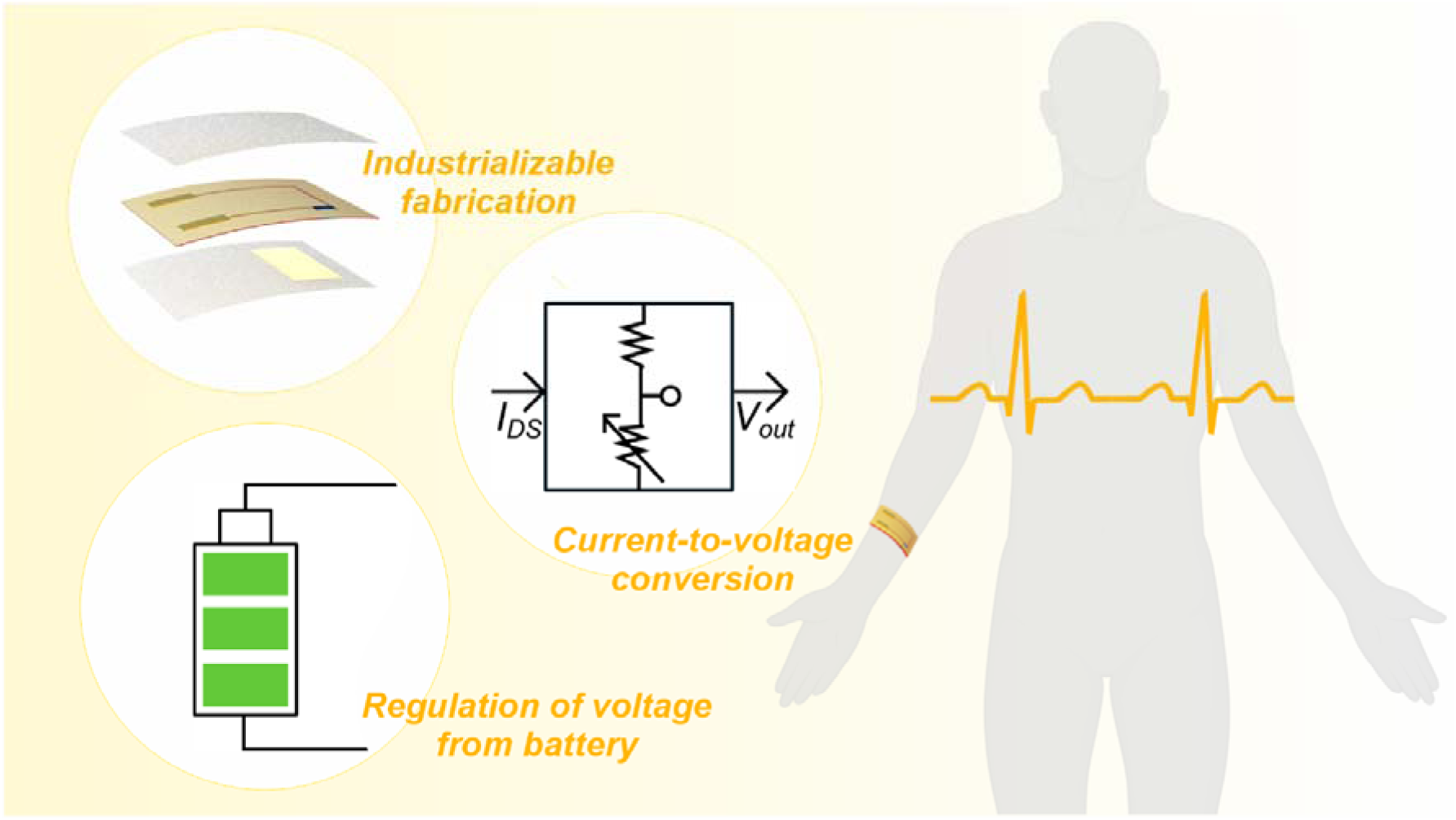
Overview of this work, encompassing the development of the fabrication flow compatible with an industrial process and the design of voltage-regulating and signal acquisition circuits for wearable OECT measurements.

## 2. Results and Discussion

### 2.1. Fabrication of Flexible OECT through Customizing Flexible Printed Circuits (FPC) Manufacturing Process

OECTs fabricated through FPC manufacturing process are composed of materials similar to the common flexible circuitry (**Figure 2A**). First, the substrate and insulating layer are made of polyimide (PI), a widely used material in microelectronics and increasingly utilized for *in vitro* and *in vivo* applications due to its excellent flexibility and biocompatibility.^[49,50]^ There is also the third polyimide-based adhesive serving as a sacrificial layer for peel-off patterning of PEDOT:PSS to be coated subsequently. The difference between FPC process and microfabrication is its relatively thick metal interconnects (i.e. 10-20 µm versus 50-100 nm), which is identified as the key factor determining whether the FPC process could be successfully adopted for OECT fabrication. Because of the absence of metal underneath the middle part of the passivation layer, the substrate at that area was exposed to laser cutting and therefore resulting in a deeper cavity between the drain and source terminals (Figure 2B). As such, the channel architecture was re-designed to a two-window configuration to accommodate the increased depth of metal leads especially for OECTs with shorter channels (Figure 2C). The dimensional metrics of both designs can be found in Figure S1.

**Figure 2.**
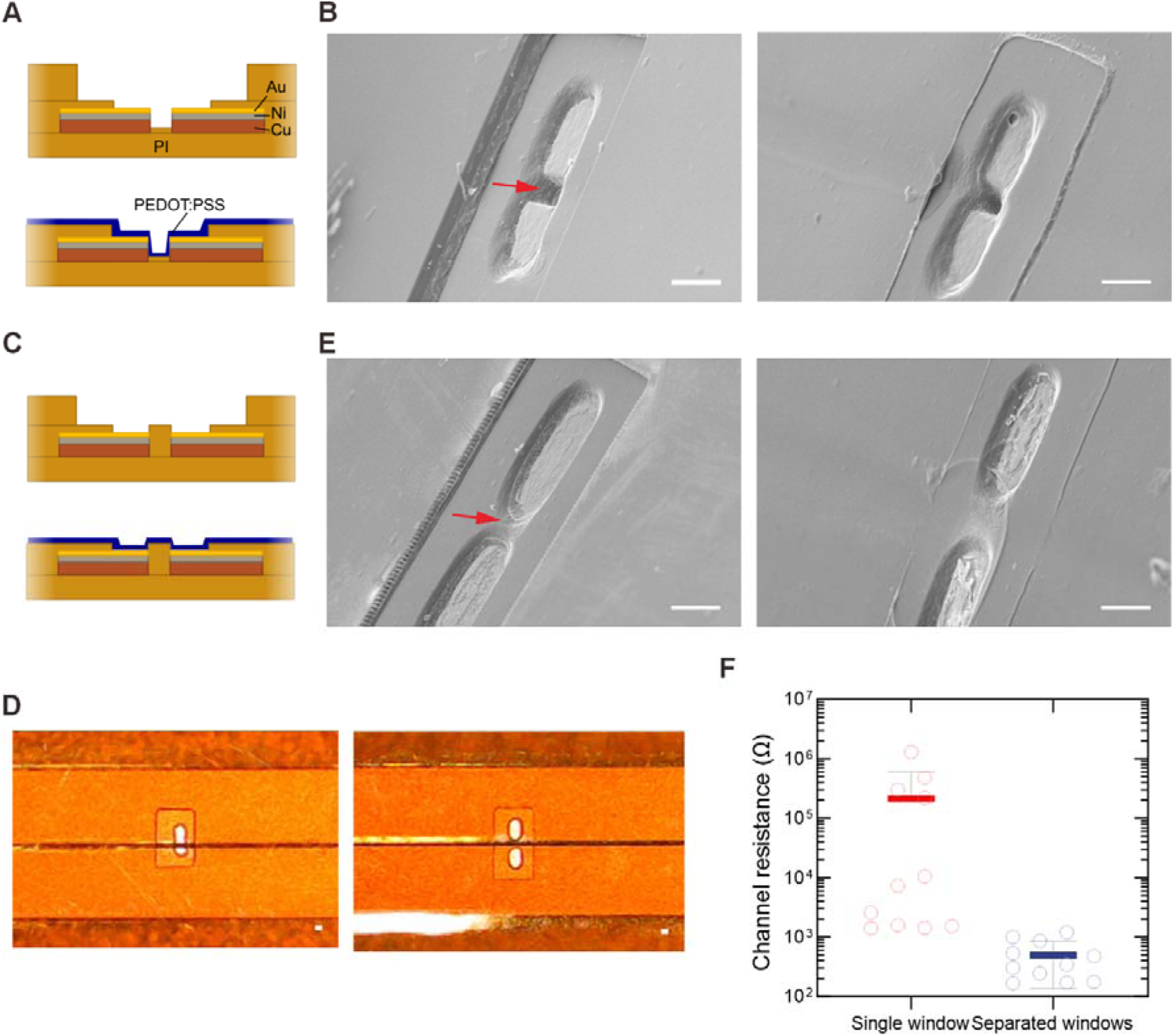
Schematic cross-sectional illustration of OECT channels with (A) single or (C) separated windows, before and after PEDOT:PSS patterning. (B) and (E) Scanning electron microscopy (SEM) images showing the top view of OECTs with configurations corresponding to the illustration in (A) and (C) respectively. Devices without (left column) or with PEDOT:PSS coating (right column). The arrows indicate the depths between the terminals and the substrate or passivation. Scale bar, 100 µm. (D) Optical images displaying transistor channels with single (left) and separated passivation openings (right) before PEDOT:PSS coating. Scale bar, 100 µm. (F) Distribution of channel resistance (mean ± SE) measured for PEDOT:PSS-coated OECTs in two different configurations. Each circle represents the resistance of one OECT. Mean values are indicated by bold horizontal lines. N = 11 transistors.

Compared to a single opening of passivation, separated openings can bypass the removal of the middle passivation (Figure 2D). Such design can minimize the depth difference between the metals and the base substrate while maintaining the exposure of two terminals (Figure 2E). The possibility of PEDOT:PSS successfully spin-coated across the depths can then be increased, leading to a much lowered channel resistance for OECTs with separated windows (Figure 2F). As proven in our previous work, a lower channel resistance can lead to a higher amplitude of “ON” current of the transistor and hence a higher amplification factor (i.e. transconductance), which favors the sensitivity of measurements ^[21]^. Therefore, the two-window passivation should be incorporated into device layout design when the aforementioned fabrication method is deployed.

Apart from device architecture, our PEDOT:PSS formulation was also optimized. Spin-coating PEDOT:PSS of conventional formulation (see Materials and Methods) on the µm-thick drain-source leads can lead to cracks at the stress-concentrated areas in the PEDOT:PSS channel, for example the corners at trenches and ridges (Figure S2). The fractured polymer possessing a higher channel resistance can in turn hamper OECT performance with decreased *I_DS_* at ON state and thus the transconductance.^[21]^ To resolve this issue, we first introduced D-sorbitol, a hygroscopic agent, that is known to increase softness, stretchability, and thickness of PEDOT:PSS.^[51]^ The improved mechanical properties are anticipated to relieve the stress, resulting in a more continuous PEDOT:PSS channel and thus a relatively high yield of OECTs with lower channel resistance consistently (Figure S2 & S3). In light of the improvements, OECTs with separated windows and optimized formulation would be used in the following experiments.

### 2.2. Signal Conversion and Amplification for OECT-based Measurements

As previously mentioned, OECT outputs are in the form of current and have to be converted into voltage signals readable and processable by downstream signal transmission and processing. In our present acquisition circuit, a load resistor (*R_load_*) is added to the drain side of the OECT to form a voltage-divider for the supply voltage (*V_sup_*) (**Figure 3A**).^[13,52]^ This configuration allows the originally fixed drain-source bias to float and reflects the *I_DS_* modulated by the biopotential (*V_bio_*) as the output voltage (*V_out_*). Despite the simplicity of this circuit, there are important points to note so as to fully harness its advantage.

**Figure 3.**
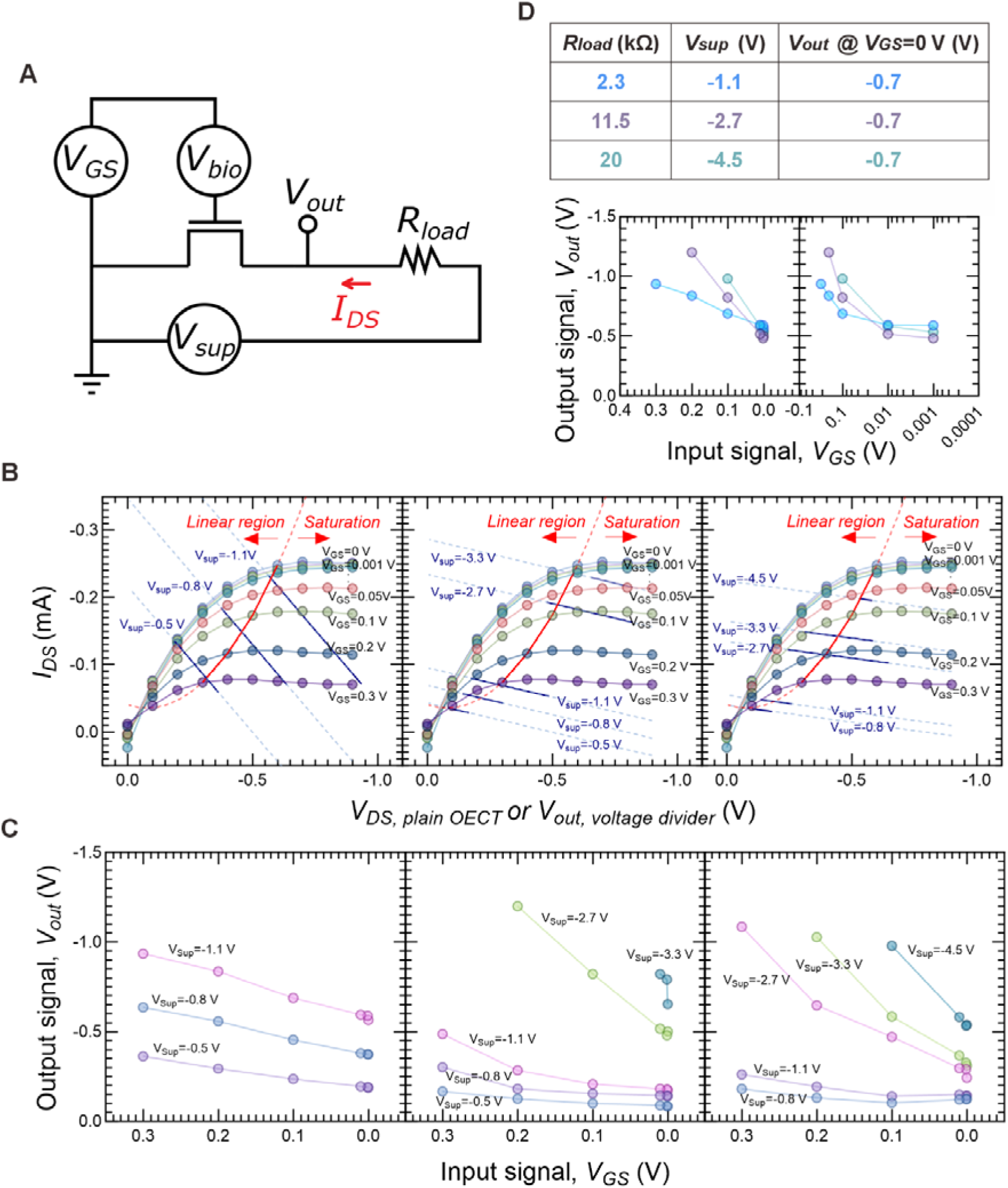
(A) Wiring diagram of the voltage-divider circuit applicable for OECT-based biological measurements, including electrophysiological recordings. Additional *V_GS_*can be applied on top of the signal of interest to bias the transistor to operate within its saturation regime. (B) Output curves (colored lines with circles) overlapped with the straight load lines (blue) for the same OECT characterized in series with a 2.3-(left), 11.5-(middle), or 20 kΩ-resistor (right). The resistance values were chosen to follow the ratio of 1:1, 5:1, and 8.6:1 to the channel resistance measured under the saturation regime, i.e. when *V_GS_*=0 V, *V_DS_*=-0.6 V. The channel resistance measured under different applied voltages can be referred to Figure S4.

Equation 1 describes the relationship of *V_out_* with other variables in the current circuit.

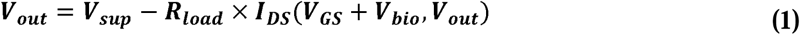

According to Equation 1, *I_DS_*amplified from the bio-signals would be further enlarged by a factor of *R_load_*to yield *V_out_*. However, *R_load_* cannot be elevated infinitely as it also draws a portion of *V_sup_*, possibly leading to an effective drain-source voltage even lower than the saturation threshold *V_DS, sat_*. Therefore, a comprehensive characterization should be performed to ensure good amplification while maintaining enough drain-source bias. In other words, the optimal values for *V_sup_* and *R_load_* have to be extracted specifically at target *V_bio_*.

To do so, output curves obtained from OECT alone are compared with *V_out_* and the corresponding *I_DS_* at the given *V_sup_* during the characterization of transistor in series with load resistor (Figure 3B). To avoid confusion, *V_DS, plain OECT_* denotes the applied drain-source voltage during I-V characterization with OECT only, whereas *V_out, voltage divider_* (as equivalent to *V_out_*) denotes the effective drain-source voltage or output voltage measured in the voltage-divider circuit. First, it can observed that the applied *V_sup_* does not necessarily lead to the exact *I_DS_* at the same *V_DS, plain OECT_*, and *V_out, voltage divider_* changes along with *V_GS_*. Taking the leftmost plot as an example, having *V_DS, plain OECT_*=-0.8 V allows only the unloaded OECT to enter the saturation region for all *V_GS_* shown. However, if the same voltage is given to *V_sup_* of a loaded OECT, its operation would remain within the linear region unless *V_GS_* exceeds 0.2 V. The latter case may require an offset *V_GS_* especially on top of the small bio-signals (e.g. ECG at around 1 mV) to achieve efficient amplification. Otherwise, the minimal required *V_GS_*(or *V_bio_*) can be decreased by providing a larger *V_sup_*, as proven grahically by the parallel translation of the corresponding load line closer to the saturation region (i.e. *V_sup_*=-1.1 versus - 0.8 V).

The red line separates the linear and saturation working regimes, and was approximated using a reported equation.^[14]^ The dotted portions are the extension from the measured data as a result of line fitting. Note that what is applied as *V_sup_* does not represent the exact drain-source voltage experienced by the transistor, and it influences *V_out, voltage divider_* (i.e. *V_out_*) together with *R_load_* and the applied *V_GS_*. (C) Graphs of the measured *V_out_* against the input voltage *V_GS_* at different *V_sup_* for the OECT connected with the 2.3-(left), 11.5-(middle), or 20 kΩ-resistor (right). (D) Enlarged linear and log plots of *V_out_*against a range of applied *V_GS_*. Data in each colored line was taken under the corresponding condition labeled in the same color in the table.

OECTs loaded with a resistance larger than its channel resistance at saturation were also tested (Figure 3B). The decrement in the slope of the load lines across the plots suggests that the magnitude of amplification increases as *R_load_* increases, which is in line with Equation 1. Meanwhile, circuits with larger *R_load_* demand larger *V_sup_* to achieve saturation. For example, OECT loaded with 2.3, 11.5, or 20 kΩ, that leads to ratios of 1:1, 5:1, and 8.6:1 to its channel resistance, would theoretically need a *V_sup_* of −1.1, −3.3, and −4.5 V respectively to saturate when the target *V_GS_* is 50 mV.

The effect of *V_sup_* and *R_load_* on amplification gain across the range of input *V_GS_* can be better visualized by plotting the measured *V_out, voltage divider_* against *V_GS_* (Figure 3C). In general for all 3 plots, providing a larger *V_sup_* results in a steeper line and thus a greater amplification. In addition, the slope of the line appears to be larger as *V_GS_* approaches to 0 V, meaning that the current device is more suitable for transduction of signals near the relevant voltage range. As examples of electrophysiological signals, ECG generally has amplitudes at 1 mV, while EMG has a broader range from 0-10 mV.^[7,53,54]^ For biochemical signals, a potential shift arising from DNA hybridization or protein-aptamer binding is reported to be 10-50 and 0-200 mV depending on concentrations, respectively.^[55,56]^

It should be noted that the gate voltage range covering the highest gain of the voltage-divider circuit also depends on the characteristics of the OECT itself, and therefore is not necessarily close to 0 V like the current case. This can be explained by the range of *V_GS_* where the maximal transconductance (*g_m, max_*) appears. In the output curves, *I_DS_* at *V_DS, plain OECT_*=-0.6 V has a relatively longer vertical distance between *V_GS_*=0-0.1 V than that between *V_GS_*=0.1-0.2 V or 0.2-0.3 V (Figure 3B, left). Similar to what is observed in the respective transfer and transconductance curves, the current OECT demonstrates a *g_m, max_*approximately at 0 V (Figure S5). As such, any load line lying across the longer vertical distance of *I_DS_* would be longer at that range of *V_GS_*, thus translating into a relatively great amplification as indicated by Figure 3C. Additional reference guiding the means to manipulate *g_m, max_* at zero gate bias can be referred to the published literature.^[57]^

In addition, lines representing larger *R_load_* in Figure 3C are distinctly steeper when *V_sup_* is provided sufficiently to saturate the OECT. For a clearer comparison on the the amplifiying characteristics loaded with different resistance, the plots conditioned at optimal *V_sup_* are extracted to Figure 3D. When *R_load_* is increased to 20 kΩ, its amplification becomes the greatest throughout the range of *V_GS_* shown (Figure 3D, left). A closer look at the range between 1 mV and 100 mV can be observed by plotting the same data in log scale, which shows a similar trend (Figure 3D, right).

To validate the above characteristic features of the circuit, we measured *V_out,_* (i.e. *V_out, voltage divider_*) subjected to a sinusoidal gate bias while alternating *V_sup_* and/or *R_load_*. As anticipated, the amplitude of *V_out_* increases as *V_sup_* is gradually tuned from −0.5 to −2.7 V and OECT becomes saturated (**Figure 4A**). Given that the OECT is saturated, *V_out_* also increases when *R_load_*is changed from 2.3 to 11.5 and 20 kΩ (Figure 4B). It should be further noted that the input gate voltage at 50 mV is amplified to at least 3 to 10 times higher depending on *R_load_*. Additionally, the amplification performance is found on par with that provided by our previous proposed transimpedance amplifier (TIA) circuit, but carrying more advantages such as lower voltage requirements and ciricuit simplicity suitable for wearable design (Figure S6 & Table S7).^[44]^ All in all, the above findings further confirm the proportionality of *V_sup_* and *R_load_* to the gain of the voltage-divider circuit, and revealed the necessary criteria to ensure satisfactory amplification of the target bio-signals such as ECG.

**Figure 4.**
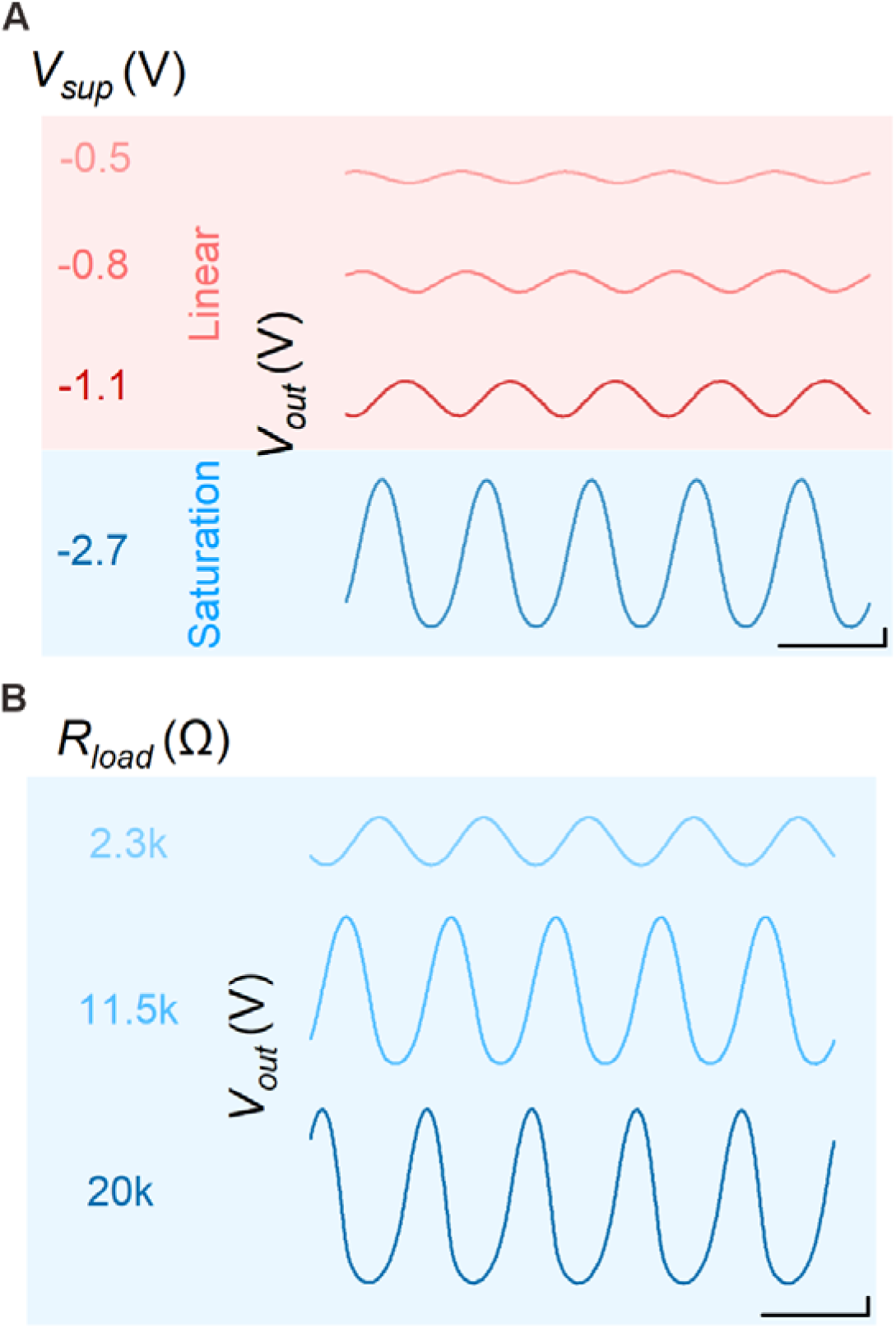
*V_out_* (i.e. *V_out, voltage divider_*) measured when the OECT is subjected to a sinusoidal *V_GS_* of *V_pp_*=50 mV at 1 Hz, (A) biased with different *V_sup_* and connected with *R_load_*=11.5 kΩ or (B) connected with different resistances with the corresponding *V_sup_*to reach saturation region. The applied *V_sup_* is −1.1, −2.7 or −4.5 V when the OECT is loaded with 2.3, 11.5 or 20 kΩ, respectively. The voltage-divider circuit is capable of amplifying by folds compared to the original signal at 50 mV. Scale bar, 50 mV and 1 s.

### 2.3. Stable and Tunable Voltage Supply for Biasing OECT

Next, we designed another circuit for sourcing stable and adjustable *V_sup_* from a battery of a fixed voltage *V_in_*. This circuit includes a voltage-regulator chip (i.e. LM337), which can regulate negative voltage output with a pair of external resistors (*R_1_* & *R_2_*) (**Figure 5a**). Other optional components include capacitors (*C_1_* & *C_2_*) for improving stability and input fluctuations, and a dummy resistor (*R_dummy_*) for preliminary characterization.^[58]^ In general, *R_1_* is fixed to 120 Ω and *R_2_* is determined from Equation 2 for a given *V_sup_*. *V_ref_* is the internal reference voltage, preset at 1.25 V.

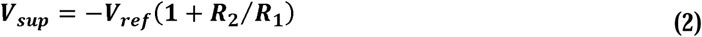

**Figure 5.**
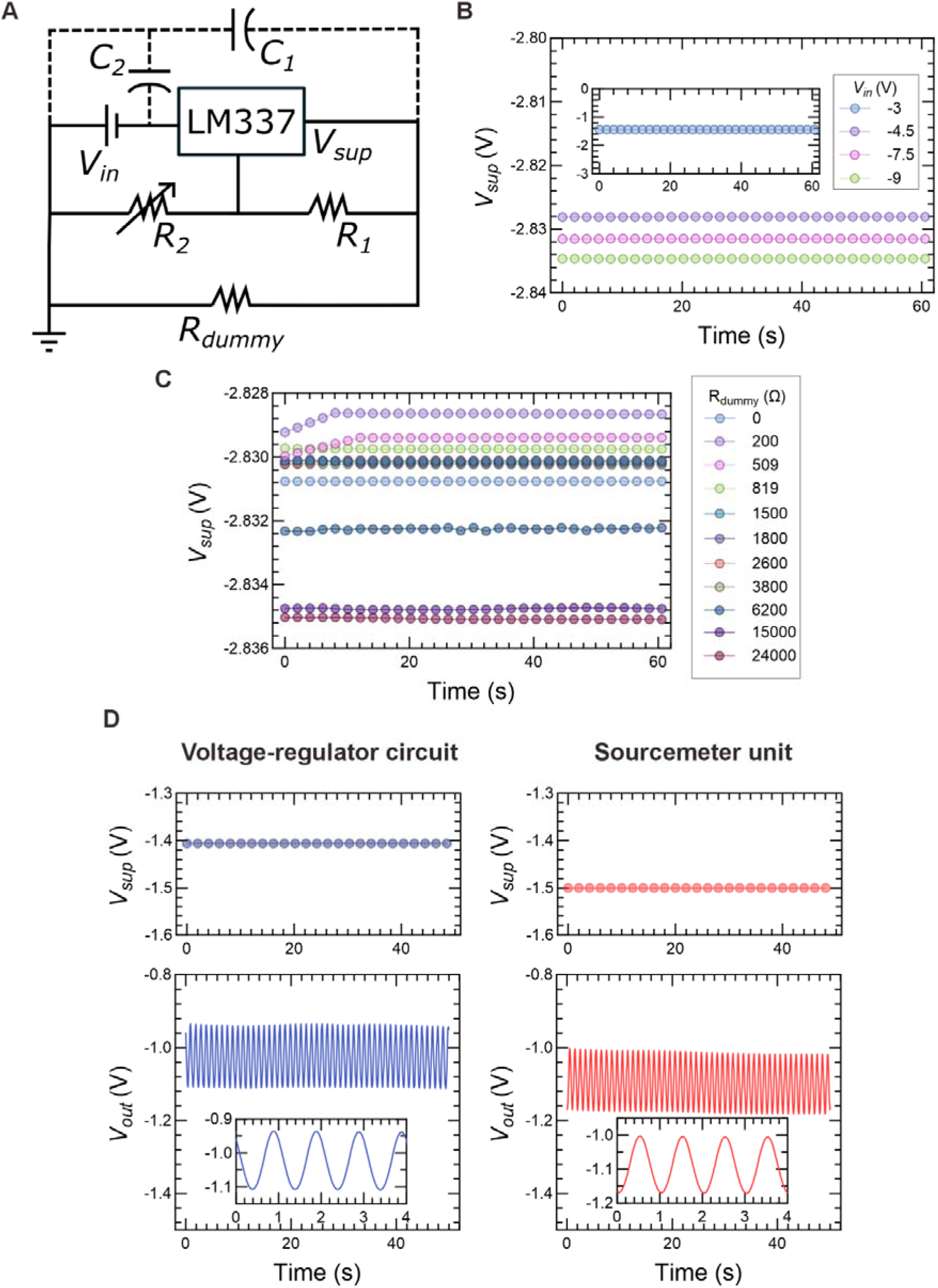
(A) Circuit diagram for supplying stable and tunable *V_sup_* from battery of a fixed *V_in_*to the acquisition circuit shown in Figure 3A. (B) Output voltage *V_sup_* over time when supplied with different input voltage *V_in_*. *R_2_* was set as 150 Ω for a *V_sup_* to be around −2.8 V. *V_in_* was provided by 1.5 V alkaline batteries connected in series. For instance, two 1.5 V batteries were used to provide *V_in_*=3 V. (C) Time stability of *V_sup_* subject to a range of *R_dummy_* representing different sourcing conditions. (D) Comparable sourcing performance between our battery-powered voltage regulator circuit and the benchtop sourcemeter unit commonly used as a constant voltage supply (top). *R_load_* was set at 24 Ω in the voltage regulator circuit to supply *V_sup_*=-1.4 V intentionally, while SMU was set to source −1.5 V directly. The corresponding *V_out_* measured from the combined circuit with OECT biased with a sinusoidal *V_GS_*at *V_pp_* of 50 mV, frequency at 1 Hz (bottom).

By substituting *V_sup_* obtained from previous characterization, *R_2_* can be determined. For example, *R_2_*would be about 200 Ω if a *V_sup_* of −3.3 V is theoretically needed for an OECT loaded with a 11.5k Ω-resistor.

The current circuit was preliminarily evaluated before integrating the two circuits together for battery-powered OECT measurements. First, we identified the minimal *V_in_* required from the battery to offer desirable *V_sup_*. This was done by decreasing *V_in_* from −9 to −3 V by removing the 1.5 V batteries in series one by one, while observing if *V_sup_* could achieve and constantly maintain a value of −2.8 V when *R_2_* was fixed to 150 Ω (Figure 5B). In our results, −4.5 V appears to be the minimum required for the circuit to supply −2.8 V. An input below −3 V can no longer maintain a −2.8 V output, despite −3 V being slightly higher than *V_sup_*. Since the attainable voltage in the current experiments is limited by the combination of 1.5 V batteries, the real minimal input/output voltage difference to achieve desirable *V_sup_*could be even smaller than suggested (i.e. in between 0.2 to 1.7 V).

In order to assess the stability of this voltage-regulating circuit, *R_dummy_* was further added in a range of values, mimicking the channel resistance under different sourcing conditions, e.g. OECT alone in the linear or saturation regions or saturated OECT with *R_load_* of ratio 1:1, 5:1, or 8.6:1 to its channel resistance. Figure 5C displays *V_sup_* over time, which varies within a negligible range (i.e. in the order of 10^-3^ V) even when *R_dummy_* is increased more than 100 folds from 0.2 to 24 kΩ (Figure 5C). All *V_sup_* measured exhibits temporal stability with no observable fluctuations and possesses limited rise time only on a few occasions.

Afterwards, this well-functioning circuit was employed to source voltage from battery to the developed voltage-divider circuit. The combined circuit was then used to capture *V_out_* as transduced from the *I_DS_* modulated by a sinusoidal gate voltage (Figure 5D, left). The stable voltage-supplying performance of this voltage-regulating circuit is shown comparable with that of the sourcemeter unit (SMU), which is a standard benchtop equipment commonly used for transistor biasing (Figure 5D, right). The sinusoidal outputs probed from the same OECT and powered separately in different settings are also clear and stable over time. Our estimation indicates that the integrated system can be operating continuously for more than 2 days before re-charging a lithium polymer (LiPo) battery with a capacity of 500 mAh (Figure S8). Together with the prior characterization, these preliminary trials substantially underscore the competence of the entire system for battery-powered ECG measurements.

### 2.4. Battery-powered Wearable ECG Measurements by OECT

**Figure 6A** illustrates the combined circuit for capturing *V_out_* of the battery-powered OECT that is modulated by biopotentials as the gate voltages. The circuit, along with the capacitors, could be assembled into a coin-sized printed circuit board (PCB) that is compact enough for wearable formats (Figure S9). Our integrated system was first used to measure heart rate signals supplied by an electrophysiological signal generator (Figure 6B). Notably, the system is capable of transducing small gate potentials (i.e. 4 mV) with amplification gain tunable by altering the load resistance. When *R_load_*is increased to 11.5 kΩ, the amplitude of *V_out_* even surpasses that measured by a conventional electrode. Simply increasing *R_load_* to a value 8.6 times larger than channel resistance can already double the bio-signals. Similar amplifying characteristics were observed in the measurement trials for lead II ECG signals (i.e. the potential difference subtracted between the voltages at left leg and right arm) supplied from the generator (Figure 6C). As shown in the data, PQRST complex, the standard characteristic pattern of electrical potentials arising from a heartbeat, can be more clearly visualized in the OECT system loaded with a 20 kΩ-resistor in comparison to the recordings by electrode.

**Figure 6.**
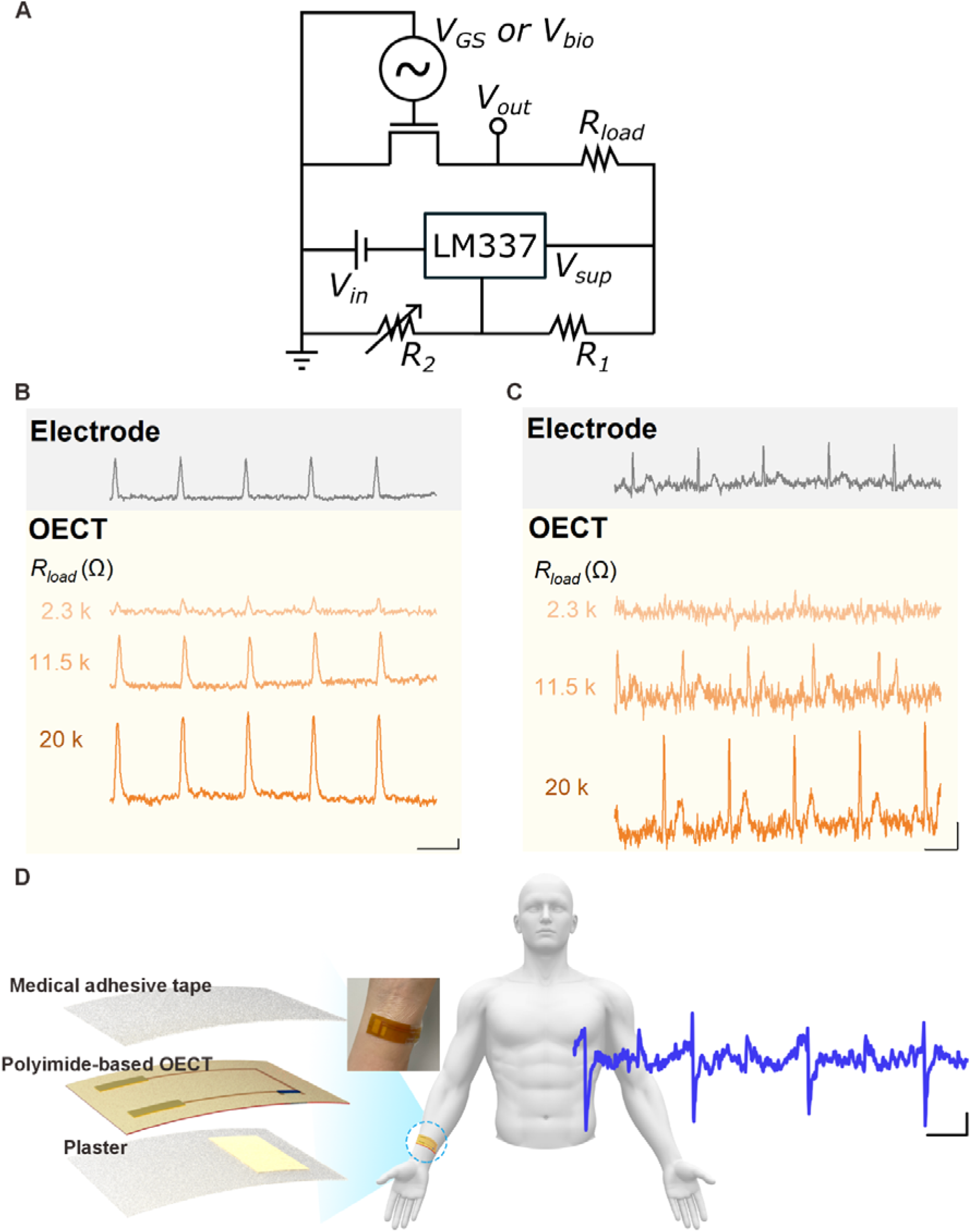
(A) The layout of the circuit with voltage-regulating and signal-acquiring functions for battery-powered OECT-based sensing. OECT can be gated by applied *V_GS_* offset, external biopotential *V_bio_*, or both. (B) Simulated heart rate signals and (C) lead-II ECG potential measured using conventional electrode or OECT integrated into the above circuit with 3 different values of *R_load_*. Scale bar, 0.5 mV and 0.5 s. In the case of *R_load_*=11.5 and 20 k Ω, *V_in_*is supplied by a 9 V battery to yield a *V_sup_* of −1.4 and −2 V respectively. In the case of *R_load_*=2.3 Ω, where *V_sup_* requires only 1 V, sourcemeter was instead used as the power source due to the limitation of minimal *V_sup_* attainable by LM337. *V_sup_* of −1.4 or −2 V was achieved by a *R_2_* of 24 or 56 Ω respectively. (D) Configuration of our flexible OECT for wearable lead-I ECG measurements (left, not drawn to scale) and the resulting ECG signals (right). Inset is the photograph of a flexible OECT mounted on skin. Scale bar, 0.5 mV and 0.5 s. In the battery-powered on-body ECG measurement, *R_load_*, *R_1_*, and *R_2_* are 20 k, 120, and 360 Ω respectively.

Our developed system was also applied for on-body ECG recordings to showcase its applicability in supporting wearable OECT-based measurements. In our prototypical device, the polyimide-based OECT is sandwiched between an ultrathin medical adhesive and a plaster, which cotton pad is soaked with liquid electrolytes to provide ions for OECT (de)doping (Figure 6D, left). The flexibility and adhesiveness of our device enable its conformal attachment to the curved surface of subject’s wrist (Figure 6D, inset). More importantly, the battery-powered OECT integrated with the developed circuits can successfully acquire the ECG potentials originated from the human subject (Figure 6D, right).

Since the same system has already been proven capable of acquiring clear ECG features from simulated signals, the relatively lower resolution shown in the measurement on the human subject could be attributed to factors relating to OECT interfaces. For instance, the ECG potentials might be partially attenuated by the impedance arising from the insulative cotton pad containing electrolytes. Also, solvent evaporation might lead to ion loss and thus ineffective (de)doping.^[31]^ Replacing liquid electrolytes with non-volatile gel electrolytes or hydrogel is one possible solution to minimize contact impedance and avoid loss of electrolytes.^[31,59]^ Alternatively, OECT with polymeric channel embedded with internal ion reservoir (also known as internal ion-gated organic electrochemical transistor, IGT) may bypass the need of external electrolytes, thus enabling direct sensing through the device itself.^[21,26,51]^

## 3. Conclusion

In this work, we presented an easy-to-build integrated module capable of regulating cell voltage for constant transistor bias while providing tunable amplification of small bio-signals transduced by FPC-like OECT. The fabrication of flexible polyimide-based OECT through the industrial FPC production process is enabled by incorporating PEDOT:PSS with D-sorbitol and patterning separated passivation windows over drain-source electrodes. The optimized OECTs demonstrate better functionality by exhibiting an averaged channel resistance 3 orders lower than that of a conventional single-window configuration. Further, the thorough and systematic characterizations of our integrated unit reveal the critical yet straightforward design rules for achieving optimal amplification of OECT’s current responses. Our proposed combination of voltage-divider and voltage-regulating circuits requires only attention to the balance between load resistance *R_load_*and supply voltage *V_sup_* controlled by another variable resistance *R_2_*. In brief, increasing *R_load_* with sufficient *V_sup_* to saturate OECT indicates elevated outputs as amplified from gate voltages at target magnitudes, and *V_sup_* supplied through a slightly higher battery voltage *V_in_* can be easily regulated by *R_2_*. Preliminary results show that our unit is able to bias OECT as stable as the standard SMU equipment and amplify into voltage outputs of magnitude 10 times higher than the input signals only by arranging *R_load_* and *V_sup_* as 20 kΩ and −4.5 V. Finally, the flexible OECT integrated with the combined circuit was successfully utilized to measure both the simulated and on-body ECG signals, with the former trials demonstrating higher quality measurements compared to electrodes. The proposed platform can be integrated not only with the current device, but also with other reported and future OECT interfaces with distinctive features, together to foster the realization of truly wearable OECTs as next-generation wearable electronics.

## 4. Materials and Methods

### 4.1. Materials

PEDOT:PSS Clevios PH1000 was procured from Heraeus. D-sorbitol, ethylene glycol (EG), 4-Dodecylbenzenesulfonic acid (DBSA), (3-Glycidyloxypropyl) trimethoxysilane (GLYMO), and sodium chloride (NaCl) from Sigma Aldrich were used.

### 4.2. Device Fabrication

All OECTs utilized in the current work were fabricated through a manufacturing process that closely resembles FPC production. First, the substrate, insulating layer, and sacrificial layer are all made of flexible polyimide. The thickness of these polyimide layers is 100, 27.5, and 20 µm respectively. The contact leads and interconnects are composed of 3 metal layers in the ascending order of copper, nickel, and gold. The metals were deposited through a well-established process called electroless nickel immersion gold process (ENIGP). The gold surface was validated by energy-dispersive X-ray analysis (EDX) (Figure S10). A mixture of conducting polymer formulated with PH1000 (0.75 g), D-sorbitol (0.2 g), EG (50 µl), DBSA (4µl), and GLYMO (10 µl) was then spin-coated on the channel at 500 rpm for 1 min, or otherwise stated. This is our improved composition of PEDOT:PSS. For conventional formulation, PH1000 (0.93 g) was mixed with EG (54 µl), DBSA (2.6 µl), and GLYMO (10 µl). The adhesive sacrificial layer with a via slightly larger than the passivation window was subsequently peeled off after a 5-min thermal annealing process at around 85 °C to pattern the transistor channel. Afterwards, the devices were further annealed at 140 °C for an hour. The PCB layout encompassing the circuit for both signal conversion and voltage regulation was drawn using the software Altium Designer. PCB boards were manufactured by Shenzhen JLC Electronics.

### 4.3. Microscopical Investigation

Optical images of the device channels were obtained using Boshile digital microscope DM4. The tilted SEM images of OECTs were acquired using the JSM-6060 scanning electron microscope from JEOL. The SEM parameters of acceleration voltage and spot size were set at 20 kV and 30 units respectively. EDX was performed alongside SEM to determine the elemental composition of the surface and thus validate the gold plating process. The analysis was conducted using the Oxford Instruments model 7582.

### 4.4. Electrical Characterization

Channel resistance was measured by Fluke 28II true RMS multimeter. I-V characterization was performed using Keithley 2612B sourcemeter (SMU) of Tektronix. Tektronix AFG31000 function generator was used to provide gate potentials in sinusoidal form. All voltage output signals, including *V_out_*, were captured either by Fluke 28II true RMS multimeter or Keysight MSO7034B oscilloscope. *V_sup_*was supplied by either GP AA batteries or SMU as stated in the main text and monitored by Keithley 2002 digit multimeter where necessary. The whole circuit for voltage regulation and signal conversion was powered by Keysight E36312A DC power supply only when measuring current for the estimation of power consumption. In all characterizing tests, OECTs were immersed in a 0.1 M NaCl solution and gated via a silver chloride-coated silver (Ag/AgCl) electrode (Shanghai Yueci Electronic Technology).

### 4.5. Simulated and on-body ECG Measurements

In the simulated measurements, voltage potentials resembling lead-II ECG pulses or heart rate signals were applied to the OECT as gate voltage. Both types of signals were generated by an electrophysiological signal generator (SKX-2000G, Xuzhou Mingsheng Electronic Technology). The simulated ECG potentials had a peak amplitude of 1 mV and a pulse rate of 60 beats per minute (bpm), while heart rate signals had a peak amplitude of 4 mV and the same frequency. The measurements of simulated signals by OECTs were configured in a setting the same as the above. In measurements by electrodes, the signals applied to the electrolytic solution via Ag/AgCl electrode were captured by the shorted drain-source terminals of the same device resembling a PEDOT:PSS-coated electrode. For the trials involving a human subject, written consent from the volunteer was obtained with the experimental protocol approved by the University’s Human Research Ethics Committee (protocol number: HREP-2022-0207). During measurements, the flexible OECT with a plaster soaked with 0.7 M NaCl solution was fixed on the subject’s right wrist by an ultrathin medical adhesive tape (YUKI-BAN PERME-ROLL Lite, Nitto). Subject’s left wrist was connected to the ground to close the circuit loop.

### 4.6. Statistical Analysis

All data in the main text and supporting information is presented without pre-processing, except Figure 6D in which the data is digitally processed via the software MATLAB using a custom code. Data presentation details are specified in the corresponding figure captions, where applicable.

## Supporting Information

Supporting Information is available from BioRxiv or from the corresponding author.

## Acknowledgements

The authors acknowledge the funding support from the Research Grants Council of Hong Kong SAR Government, China (GRF 16302723). S. Y. Yeung would like to thank the HKSAR government for providing Hong Kong Ph.D. Fellowship and Electronic Packaging Laboratory (EPACK Lab) of HKUST for technical support. Technical advice from Mr K. C. Peter Lee is also recognized.

## Data Availability Statement

The data that support the findings of this study are available from the corresponding author upon reasonable request.

