## Supplementary information for "Battery-powered Wearable Utilizing Flexible Printed Circuit-based Organic Electrochemical Transistor Embedded with Simple Circuits of Voltage Divider and Regulator for Biosignal Measurement"

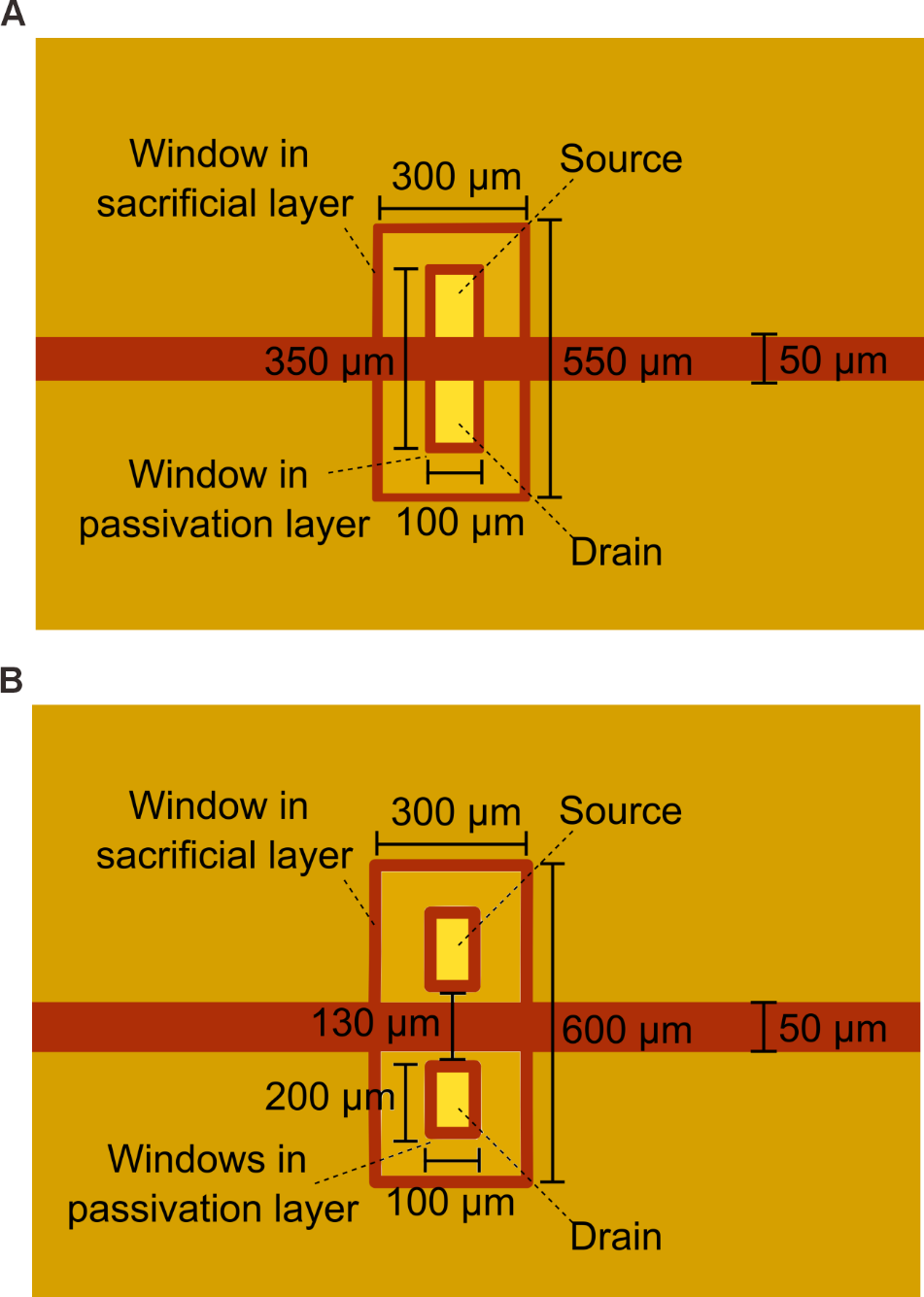

**Figure S1**. Layout of OECT channels with (A) single or (B) two windows before PEDOT:PSS coating. Not drawn to scale.

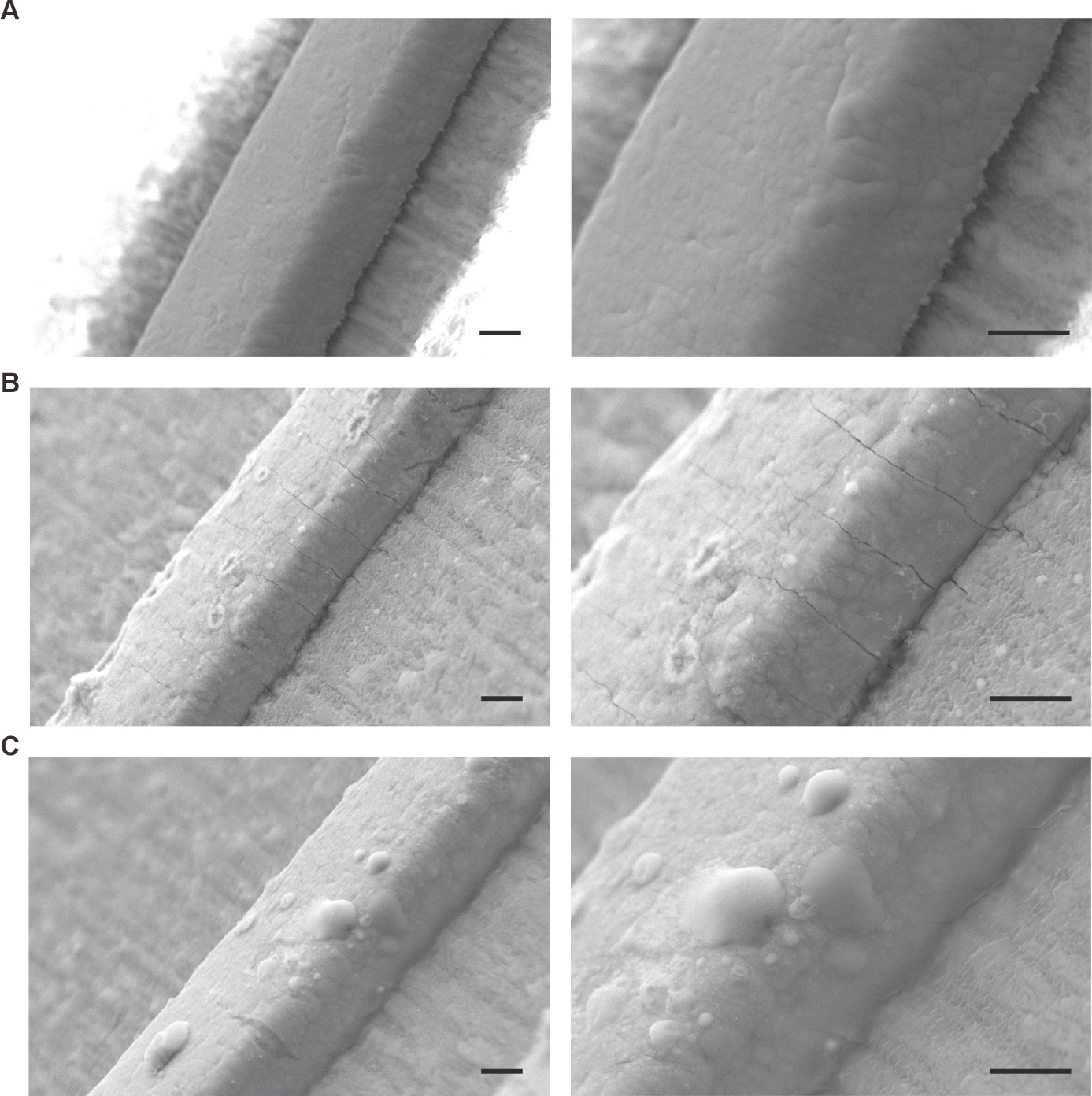

Figure S2. SEM micrographs of polyimide-based OECT (A) before PEDOT:PSS coating, (B) coated with PEDOT:PSS of conventional formulation, or (C) coated with our optimized formulation with the focus on the metal leads.^[1]^ Images were taken under resolutions at 1000x (left column) and 2000x (right column). Detailed composition can be found in the Experimental Section. The devices shown the current images have a channel width and length of 3 and 1 mm, respectively. Scale bar, 10 µm (left column) and 10 µm (right column).

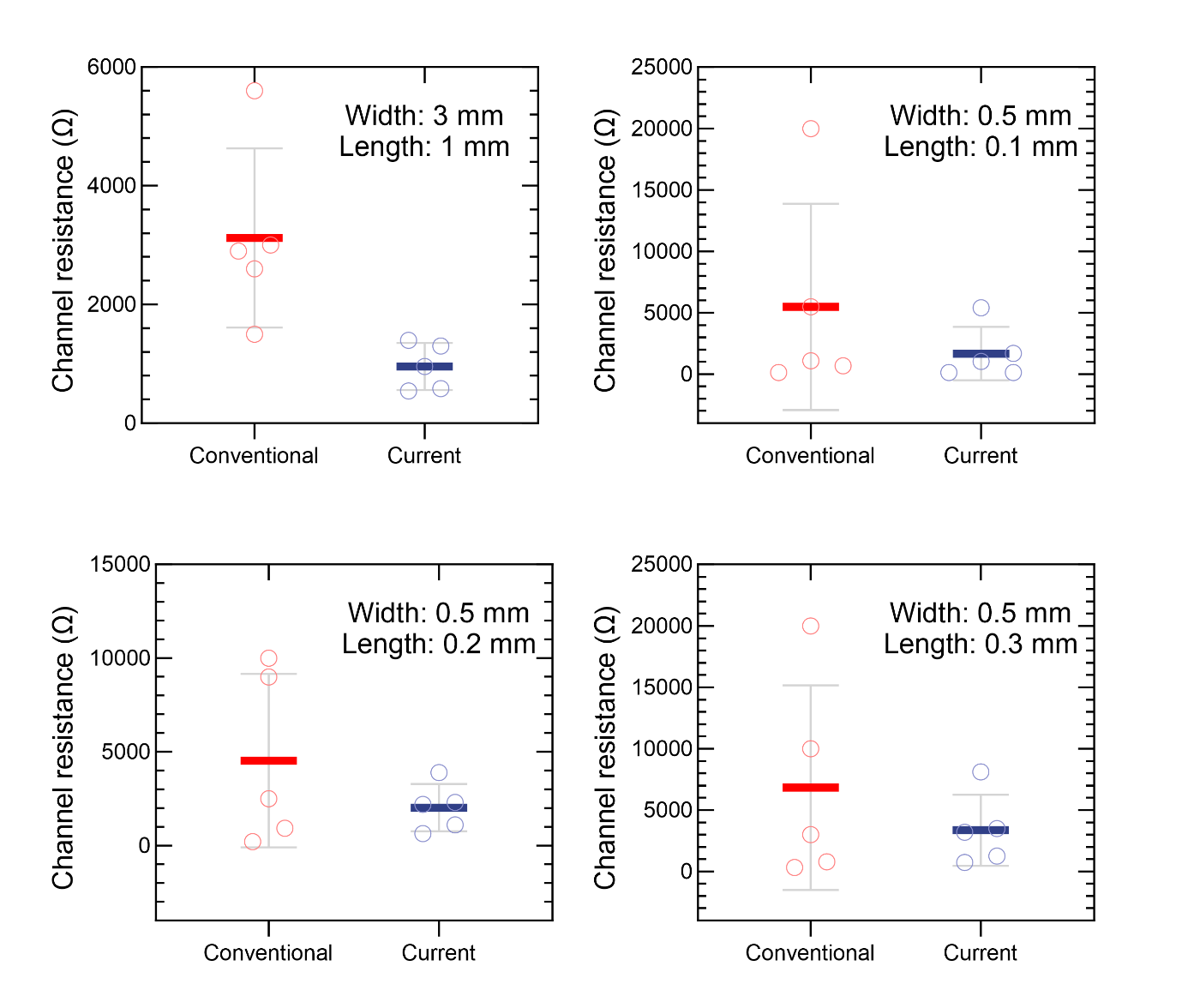

Figure S3. Distribution of channel resistance (mean ± SE) measured for OECTs coated with conventional or improved formulation. OECTs fabricated through FPC process likely have a more dispersed and higher channel resistance when coated with conventional PEDOT:PSS formulation. The improving effect of compositional changes on the averaged channel resistance is more significant when the device dimension is larger. Each circle represents the resistance of one OECT. Mean values are indicated by bold horizontal lines. n = 5 transistors for each channel dimension.

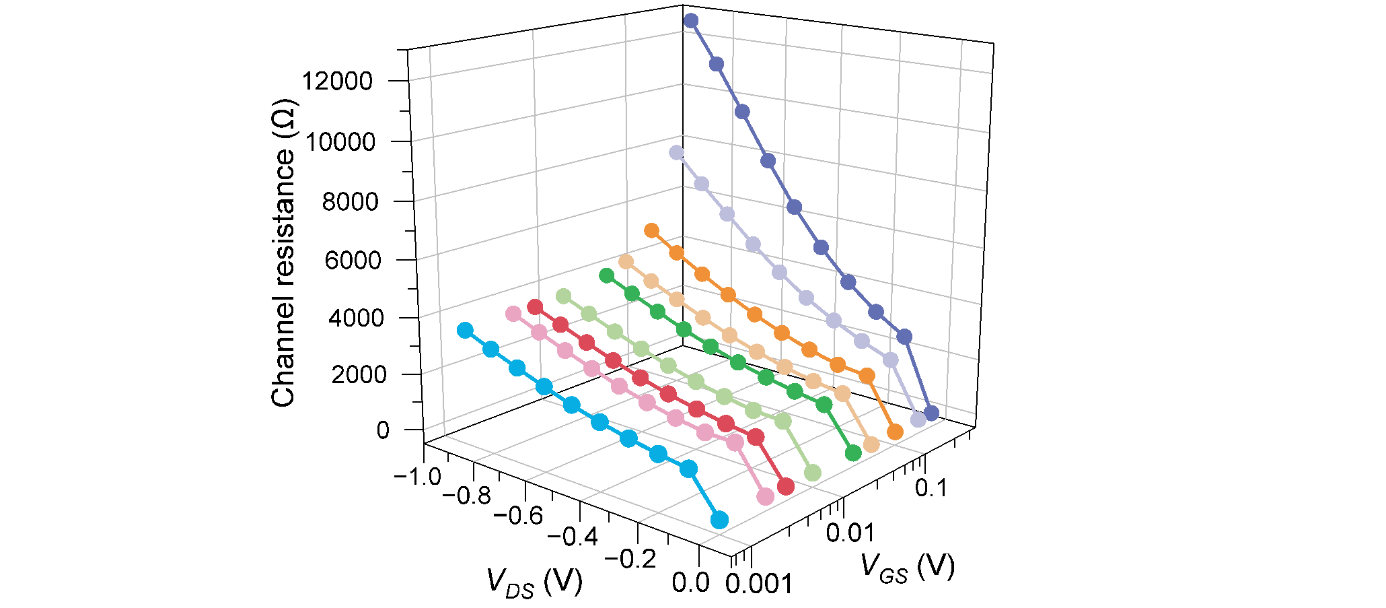

**Figure S4.** A 3D plot of OECT resistance at varying *V**_DS_* and *V_GS_*. The channel resistance at where OECT enters the saturation region was presumably taken at *V_DS_*=-0.6 V and *V_GS_*=0 V, which is approximately 2.4 kΩ. This value of channel resistance was recruited to determine *R_load_* at different ratios, i.e. 1:1, 5:1, and 8.6:1. Moreover, the channel resistance at *V_G_*_S_=0 V can resemble the resistance of OECT biased by external ECG potential, which is typically at the range of 1 mV.^[2]^

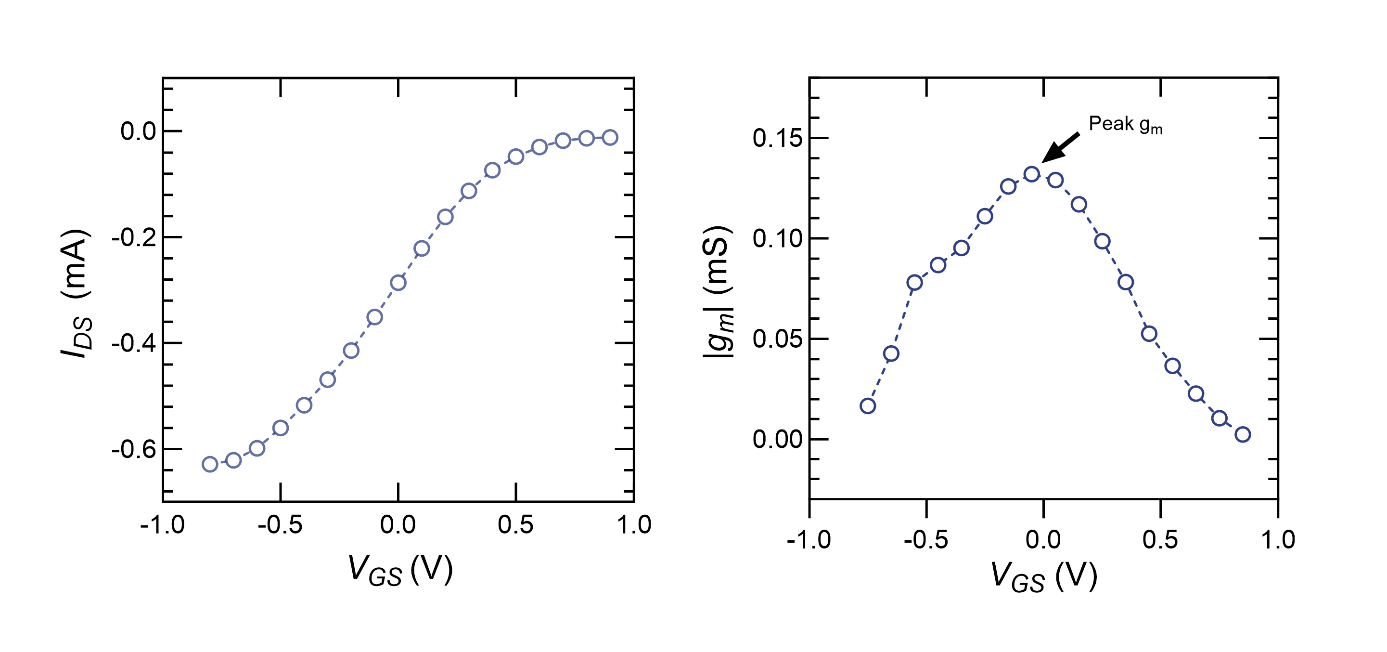

**Figure S5.** (A) Transfer curve and (B) transconductance characteristic of the OECT employed in the main text. The transconductance *g_m_* reaches its maximum when *V_GS_* approaches 0 V.

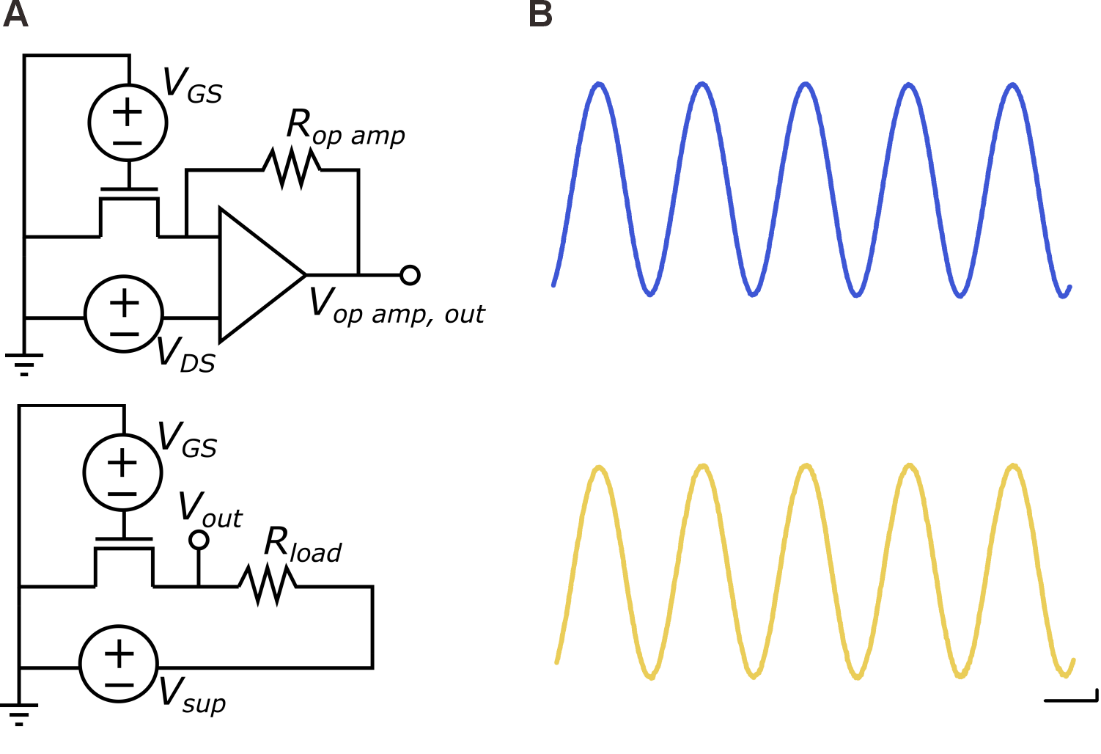

**Figure S6.** (A) Diagrams of our previously reported current-to-voltage op-amp circuit (top) and the proposed circuit in this work (bottom).^[3]^ (B) Voltage outputs obtained from the same OECT configured in the two different circuits shown correspondingly in (A). Both *R_op amp_* and *R_load_* are 10 kΩ, resulting in the same theoretical amplification factor. Scale bar, 5 mV, 0.5 s.

**Table S7.** Comparison between our reported current-to-voltage op-amp circuit and the circuit proposed in this work.

|  | Our previous current-to-voltage conversion circuit^[3]^ | This work (voltage-regulating and signal acquisition circuit) |
| --- | --- | --- |
| Voltage required for constant voltage source and signal acquisition circuit (V) | - +5 V & -5 V for -0.38 V constant voltage - +15 V & - 15 V for op-amp | - -6 V for a voltage-divider circuit designed to consume -3 V |
| Ease to use | Lots of connections | Relatively simple |
| Accuracy | Comparable | |

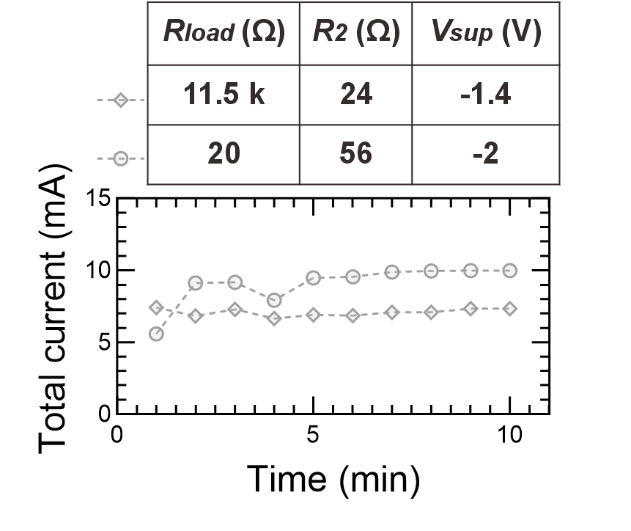

**Figure S8.** Measured current of the whole voltage-regulating and signal acquisition circuit over time.

The time duration of operation supported by a battery of given capacity can be estimated by the following equation.

$$\boldsymbol{Duration of operation per charging (h)=}\frac{\boldsymbol{Total capacity (mAh)}}{\boldsymbol{Load current}\left( \boldsymbol{mA} \right)}$$

(1)

Considering the maximal current observed in the above data (i.e. 9.98 mA), a 7.4 V, 500 mAh lithium polymer (LiPo) battery can support the operation of OECT integrated in the proposed circuit continuously for more than 50 hours, which is reasonably sufficient for use before another round of battery recharging.

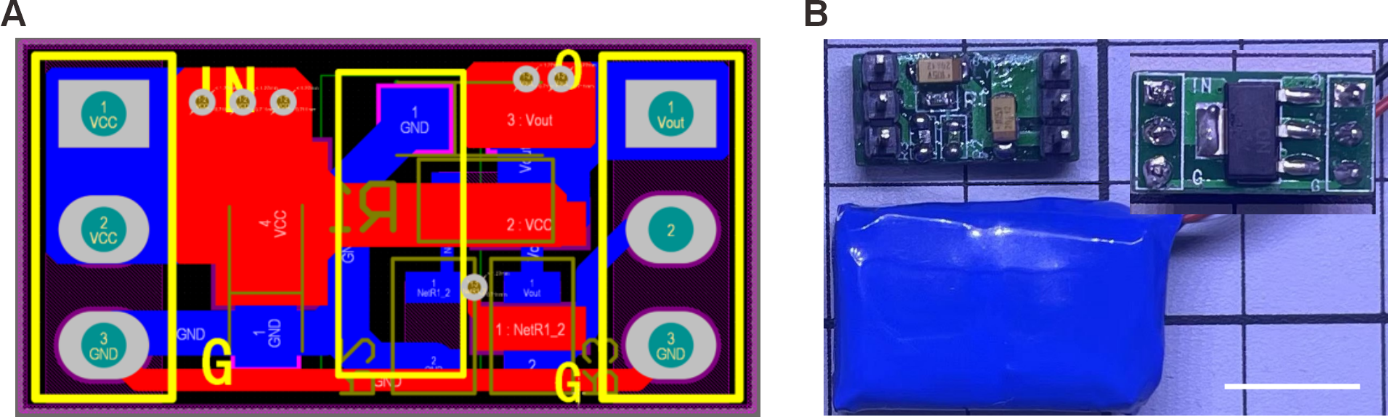

**Figure S9.** (A) Layout of the PCB designed for the proposed circuit, where capacitors *C_1_* and *C_2_* are also included. (B) Photograph of the assembled PCB and a typical 7.4 V, 500 mAh LiPo battery. Inset shows the backside of the PCB. Scale bar, 1 cm.

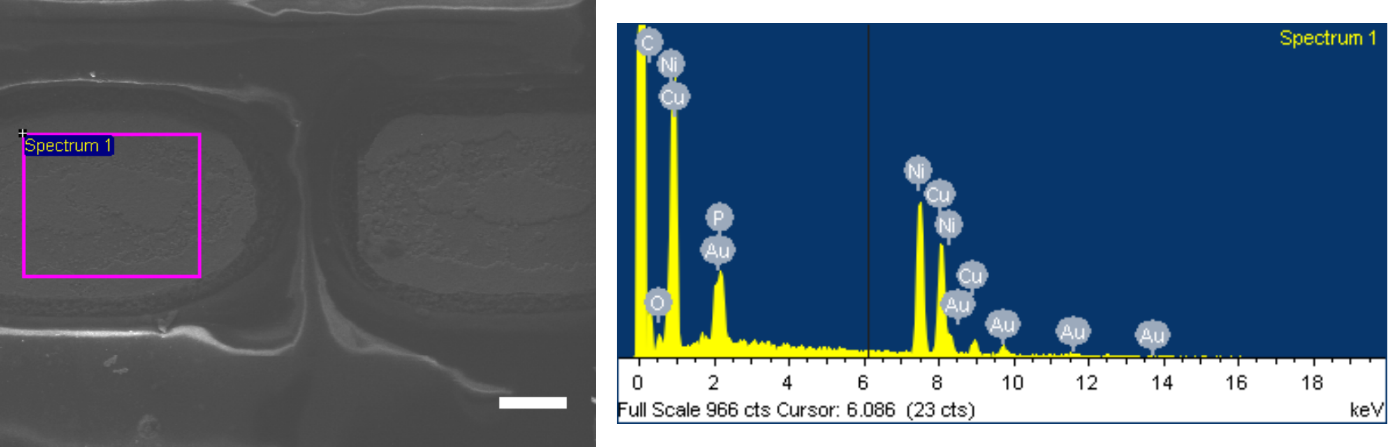

**Figure S10.** The EDX spectrum (right) revealing the types of elements automatically detected on the surface of the labelled area in SEM image (left). The area lays within the metal terminal of the fabricated OECT. Scale bar, 50 µm.
